# Anterior corpus callosum modulates symmetric forelimb movements during spontaneous food handling behavior of rats

**DOI:** 10.1101/628248

**Authors:** Masakazu Igarashi, Yumiko Akamine, Jeffery R Wickens

## Abstract

Bimanual motor actions, such as threading a needle, require coordination of the movements of each hand according to the state of the other hand. By connecting homologous cortical regions between the two cerebral hemispheres, the corpus callosum is thought to play a key role in such bimanual coordination. However, direct experimental evidence of the contribution of the corpus callosum to natural behaviors requiring bimanual coordination, such as feeding, is lacking. We investigated the hypothesis that the corpus callosum mediates bimanual movements during food-handling behavior. We first traced the forelimb-related components of the motor corpus callosum in Long-Evans rats, and found that the callosal fiber bundle from the forelimb motor areas passes through the anterior part of the corpus callosum. We then confirmed by electrophysiological recordings that blocking the axonal conduction of fibers in the anterior corpus callosum reduced neural transmission between cortical forelimb areas. The causal role of corpus callosum in bimanual coordination was then tested by analyzing forelimb kinematics during object manipulation, before and after blocking axonal conduction in the anterior corpus callosum. We found the frequency of occurrence of symmetric bimanual movements was reduced by inhibition of anterior corpus callosum. In contrast, asymmetric bimanual movement was increased. Our findings suggest that the anterior corpus callosum coordinates the direction of bimanual movement.

## Introduction

Bimanual coordination is one of the most frequently observed motor behavior in our daily life (Vega-González and Granat 2005). Human exhibit advanced capabilities for bimanual coordination. For example, controlling a steering wheel while turning a corner requires both hands. Similarly, bimanual coordination is necessary for the most basic needs of living, such as feeding, which are also important for non-human primates and rodents (Brinkman 1981; Whishaw and Coles 1996). To use both hands in a coordinated manner, motor commands between the two sides of the body must be integrated. Given that the motor representations are lateralized in primary motor cortex (Boldrey and Penfield 1937), i.e., the sides of the body are represented in separate hemispheres, a bilaterally interconnecting neural structure may be crucial for coordinating the right and the left side. The corpus callosum is a direct commissure between cerebral hemispheres, which may provide integration of motor command between contralateral cortices.

The corpus callosum is thought to be important for bimanual coordination because it provides neural connections between homologous cortical regions in the two hemispheres cortices (Hofer and Frahm 2006; Gooijers and Swinnen 2014). Such connectivity favors concurrent activation of homologous motor representations bilaterally. In support of this idea, a spontaneous transition from asymmetric motor pattern (activation of muscle groups in different timing) to symmetric motor pattern (activation of different muscle groups in same/different timing) has been reported in many experimental conditions, such as rhythmical bimanual finger tapping and bimanual drawing (Franz *et al*. 1996; Swinnen *et al*. 1998; Eliassen *et al*. 1999; Kennerley *et al*. 2002).

Rodents exhibit bimanual coordination when feeding; holding and manipulating food, and bringing the food to the mouth. However, experimental studies of bimanual coordination in rodents have been relatively limited to date. Recently, measurement techniques for bilateral forelimb movements during spontaneous food handling have been developed using high-speed camera. These techniques enable the analysis of kinematic parameters of forelimb movements bilaterally (Igarashi and Wickens 2019). Therefore, it is now possible to address the hypothesis that the corpus callosum mediates bimanual movements by quantifying symmetric forelimb movement bilaterally in time and space.

In the present study, the contribution of the corpus callosum in bimanual coordination is investigated by measuring food handling behavior in head-fixed rats. Our working hypothesis is that the motor corpus callosum is important for symmetry in movements. To address this question, we identified motor pathways in the corpus callosum that connected the two forelimb motor areas n rats, and confirmed that this motor callosal connection was temporarily silenced by a sodium channel blocker. We then examined the effect of blockade of the anterior corpus callosum on bimanual coordination by 3-D kinematic analysis of feeding behavior. Kinematic analysis was used to test two specific hypotheses (i) the anterior corpus callosum mediates symmetry in forelimb movement speed; and, ii) the anterior corpus callosum mediates symmetry in forelimb movement direction. We found that suppressing the anterior corpus callosum connections altered the ratio of movement symmetry toward more asymmetry in movement direction. However, symmetry in movement speed was unchanged. Other aspects of skilled forelimb use, such as time of consumption, were unchanged. We suggest that the anterior corpus callosum is important for integrating spatial representation of two motor cortical forelimb areas to generate fine symmetric bimanual movements.

## Materials and methods

### Animals

Thirty four Male Long Evan rats weighing 350-450 g were used in the study. Rats were kept under a reversed 12 hrs light/dark cycle, constant temperature and humidity with free access to water and food before food restriction. Animals were habituated to the experimenter for more than three days before the start of behavioral training. All experiments in the present study were approved by the Committee for Care and Use of Animals at the Okinawa Institute of Science and Technology.

### Tracing callosal fibers from forelimb motor areas

Eight rats were used to visualize the connections between the forelimb motor areas. These rats were anesthetized with isoflurane (4.0% induction, 1.5-2.0% maintenance). To visualize the axonal fiber bundle, 1,1’-Dioctadecyl-3,3,3’,3’-Tetramethylindocarbocyanine Perchlorate (DiI, Invitrogen, USA) was used as a non-viral neural tracer. 2% DiI solution was prepared by dissolving DiI crystals in 100% ethanol. 100 nL of the 2% DiI solution was then slowly injected into either rostral forelimb motor area (RFA) or caudal forelimb motor area (CFA) at 1 nl/sec (Nanoject III, Drummond Scientific). The coordinates for the RFA and CFA in male Long Evans rats were based on previous reports (Brown and Teskey 2014; Neafsey and Sievert 1982) (RFA: 3.2 mm anterior and 2.3 mm lateral from bregma, 0.8 mm deep from cortical pia; CFA: 1.0 mm anterior and 2.5 mm lateral from bregma, 0.8 mm deep from cortical pia). Animals were kept alive for two weeks to allow for the diffusion of DiI. The rats were then euthanized with pentobarbital and perfused with 100 mL of heparin saline solution followed by 100 mL of 4% PFA in Phosphate Buffer (PB). Brains were removed and post-fixed with 4% PFA for 2 hours followed by two days of saturation in 30% sucrose PB solution. The brains were sectioned at 100 μm and the slices were then rinsed in PB and nuclear DNA was stained with the NucBlue™ Fixed Cell Stain using the manufacturer’s protocol (Life Technologies, Grand Island, NY). The fluorescence of DiI and NucBlue™ were observed under a fluorescence microscope (BZ-9000, KEYENCE, Osaka, Japan) using DAPI and mCherry filters.

### In vivo anesthetized LFP recordings

Eight rats were used for recording of cortical local field potentials (LFPs). Animals were placed on a stereotaxic frame under isoflurane anesthesia (3.0% induction, 1.5% maintenance and recording). Two millimeters square craniotomies were made on both right and left sides of the skull centered on the central region of primary motor forelimb area in Long Evans rats (Neafsey and Sievert 1982; Brown and Teskey 2014) (0.5 mm anterior and 1.5mm lateral from the bregma). A 16-ch array silicone probe (A1×16-5mm-150-413, NeuroNexus, USA) was inserted to the depth of 2.2 mm from the pia to record LFPs from cortical laminae. A concentric bipolar stimulation electrode (CBARC_75, FHC Inc., ME, USA) was slowly inserted 1.0 mm from the cortical pia contralateral to the silicone probe. A glass pipette connected to a microinjector was inserted into the anterior corpus callosum for the injection of sodium channel blocker (Nanoject III, Drummond Scientific, PA, USA, 0.5 mm anterior and 0.8 mm lateral from the bregma, 2.9 mm deep from the cortical pia). A wideband signal was recorded using an OmniPlex D multichannel recording system (Plexon, TX, USA). The signal was filtered with a 200 Hz lowpass cutoff Bessel. Intracortical microstimulation (ICMS) was applied to measure cortico-cortical connections. Cathode-lead biphasic current pulses (±150 μA, 1 ms) were applied every 10 seconds using a programmable stimulus generator (STG4004, Multichannel Systems, Germany). The first 10 minutes were used for baseline. Immediately after the baseline, 500nL of 2% Lidocaine solution was injected at 1nL/sec and recordings were continued for 30 min. After completion of LFP recordings, rats were perfused with 4% PFA for histology, the brains were sectioned on a vibratome and stained with NucBlue™ or subjected to Nissl staining to show cells. Fluorogold and NucBlue™ were imaged under a fluorescence microscope (BZ-9000, KEYENCE, Osaka, Japan) using DAPI and GFP filters.

### Data analysis of LFP recording

The recorded continuous 16-ch LFP data set was segmented and realigned to the time onset of ICMS events to produce peri-stimulus LFP traces, using Neuro Explorer (Nex Technologies, MA, USA). The mean of peri-stimulus LFPs were then computed from every 10 minutes of recording. The largest sink of LFP among 16 channels was selected as the peak LFP response. To visualize current source density (CSD) profile, the mean peri-stimulus LFP traces were exported to MATLAB and inverted current source density (iCSD) plots were generated using iCSD plot toolbox for MATLAB distributed by Pettersen *et al*. (2006).

### Surgery for head-fixation

Ten to twelve week old male rats weighing 350-450g were used for behavioral experiments. Rats were anesthetized with isoflurane (3 – 4% induction, 1.5 – 2.5 % maintenance), and placed on a stereotaxic frame (SR-10R-HT, Narishige, Japan). The detailed procedures of implantation of head-plate has been previously described (Igarashi and Wickens, 2018). Briefly, the skull was exposed and carefully cleaned with saline and cotton swabs. The eight anchor screws drilled to the skull were then covered with a layer of Super Bond (Sun Medical Inc., Japan). A chamber frame (CFR-1, Narishige, Japan) was positioned and secured by additional dental cement. Dietary supplement with Carprofen (Medigel CPF; Clear H_2_O, ME., US.) was given during post-op recovery for 5 days.

### Spontaneous food handling under head-fixation

Rats were placed on the food deprivation protocol 1 week before habituation. Behavioral testing was conducted during the middle hours of the dark cycle (10am – 4 pm, reversed light cycle). At the time of testing, the last feed had been given to the animals on the previous day. Testing was conducted on one animal at a time in a quiet room. Rats were habituated to the experimental chamber for three days, and gradually guided to the head attachment clamp by the experimenter using a sweet jelly reward (Igarashi and Wickens, 2018). To elicit bimanual motor behavior, a modified version of the spontaneous food handling task originally proposed by Whishaw and Coles (1996) was used. The annular shaped food reward (20 mm outer diameter, 10 mm inner diameter, 5 mm thickness, Fish Sausage, Marudai Food Co., Ltd, Japan) was used instead of vermicelli or pasta. Trials started with bimanual grasping of the food reward offered by the experimenter, and movements of the forelimbs during consumption were recorded. A successful trial was defined as complete consumption of a single food reward without dropping it. Rats underwent 6 trials in a day and continued for 6 days (Fig. 3). Cases where rats showed unusual behavior, such as adopting a tripedal stance during eating, were excluded from further analysis.

### Intracallosal drug infusion

After training, rats were anesthetized with 2% isoflurane, placed in a stereotaxic frame, and implanted with a stainless guide cannula (26G, 7mm, PlasticsOne, VA, USA) into the anterior part of the corpus callosum (1.0 mm anterior posterior and 0.8 mm lateral from bregma; 2.0 mm ventral from cortical pia). Due to the blood vessels along the sagittal sinus, the cannula was placed 0.8 mm lateral from the midline. The cannula was fixed by dental cement (Super-Bond, Sun Medical Inc., Japan) and secured by a dummy cannula (7mm, PlasticsOne, VA, USA). A dietary supplement of Carprofen (Medigel CPF; Clear H_2_O, ME, USA) was given during post-op recovery for 5 days. 15 min prior to behavioral test sessions, 500 nl of 2% Lidocaine (dissolved in saline) was injected via an internal cannula (1.0 mm exposure from guide cannula) using a 10 μl gas tight syringe loaded on a syringe pump (KDS-101, KD Scientific Inc., MA, USA). Lidocaine was slowly injected at a rate of 1.67 nl/sec (5 min total) and the internal cannula was kept in position for 5 min to allow diffusion. After completing six sessions of behavioral experiments, rats were deeply anesthetized and perfused. The brains were sectioned and the location of the tip of cannula was confirmed by Nissl staining.

### Recording of forelimb motor behavior and 3-D reconstruction

On the day of the behavioral recording, 3 mm diameter half-spherical reflective markers were attached to the lower side of the wrists with double-sided tape. Rats were head-fixed in the custom-made apparatus (SR-10R-HT, Narishige, Japan; Fig.). The reflective markers attached to the forelimb were tracked during food handling by two high-speed cameras (HAS-L1, f = 6mm, DITECT, Tokyo, Japan) positioned below the transparent acrylic floor. All trials were recorded at 200 frames per second (1/500s exposure time and 600×800 pixel) and stored to hard disk. The positions of reflective markers were semi-automatically traced and reconstructed using custom MATLAB programs (The MathWorks, Inc., MA., USA). The positions of the reflective markers were represented as a time series data in the camera coordinate [x, y, z]^*T*^, where x = [x_1_, x_2_, …, x_t_], y = [y_1_, y_2_, …, y_t_], z = [z_1_, z_2_, …, z_t_]. The data [x, y, z]^*T*^ were transformed into the egocentric coordinate system [lr, ap, dv]^*T*^ using a reference frame based on the head-fixed apparatus (Igarashi and Wickens, 2018), where lr, ap, dv corresponds to time series data of marker position in left-right (lr) axis, anterior-posterior (ap) axis, and dorsal-ventral (dv) respectively.

### Analysis of kinematic data

To analyze organization of bilateral forelimb coordination, laterality of movement speed and asymmetry in movement direction were computed as detailed by Igarashi and Wickens (2018). The kinematic data was analyzed by following three steps:

1. *Detection of forelimb movements*. Rats demonstrated frequent transition between resting states and active use of forelimbs during food consumption. The active use of two forelimbs was detected by the maximum speed function 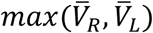, where 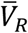 and 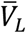 are mean speed computed from [lr, ap, dv]*^T^* by each 50 ms sliding time window. The threshold for detecting movement was set to 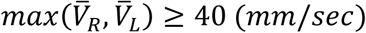, that is, if mean speed of either left or right forelimb exceeded 40 mm/sec the time frame was defined as *movement*. This threshold value was previously validated (Igarashi and Wickens, 2018). Conversely, the resting state 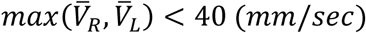 was excluded from further analysis.
2. *Speed ratio.* The extracted active movements were further analyzed by the speed ratio function 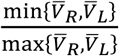. The speed ratio function was used to compute synchronization of movement speed across two forelimbs disregarding movement direction. The speed ratio function uses the speed of left and right forelimbs 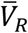 and 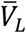 to calculate the ratio of the larger value to the smaller value (e.g. 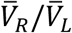 in the case of 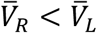). SpeedRatio = 1 indicates that the speed of both forelimbs is identical (bilateral movement), whereas a SpeedRatio = 0 suggests that the speed of one forelimb is zero (unilateral movement). Bilateral and unilateral forelimb movements were detected by setting the threshold to 0.5.
3. *Asymmetry index.* Asymmetry in movement direction was computed using an inverse cosine similarity function 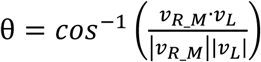, where theta is a measure of the angle between the movement vectors of the left forelimb *υ_L_* and the mirrored right forelimb *υ_R_M_*, which was obtained by mirror transformation of movement vector of the right forelimb *υ*_R_ with respect to the sagittal plane. The mean asymmetry index 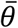 was computed in each 50 ms sliding time window. For quantitative analysis of the asymmetry index during behavioral testing, a threshold value of π/4 (45 degree) was used to classify movements into one of two categories: i) symmetric movements, ii) asymmetric movements. Orthogonal lever pressing with two hands has previously been used as an example of asymmetric bimanual movements, which is neurophysiologically significantly different from perfect symmetry (0 degree) (Cardoso de Oliveira *et al*. 2001). The present study used the value π/4 which is intermediate between perfectly symmetric movement (0 degree) and orthogonally asymmetric movements (90 degree).

Global scores of spontaneous food handling behavior were defined as follows: Mean speed of forelimbs was defined by the grand mean of movement speed in each trial. The rate of successful food consumption was calculated by the number of failed trials (a drop of food) divided by the total number of trials. The mean time of completion of food intake was the grand mean of the time spent on the consumption of single annular shaped food reward. The cross correlation between forelimbs was calculated from the cross-correlation of movement velocities between mirrored right forelimbs *υ_R_M_* and the left forelimb *υ_L_*.

## Results

### Rat forelimb cortical motor areas project in anterior corpus callosum

Previous work in humans has shown that the corpus callosum is functionally organized along the anteroposterior axis with respect to the origin of cortical areas (Doron and Gazzaniga 2008). In the present study the callosal bundle from the motor forelimb areas in rats was localized by injecting the neural tracer DiI into two forelimb motor areas: the rostral forelimb area (RFA) and the caudal forelimb area (CFA) (Fig. 1A). DiI positive axonal fibers could be clearly identified in their course through the corpus callosum (Fig. 1B). DiI positive fiber bundles from both CFA and RFA coursed mainly in the anterior part of the corpus callosum (Fig. 1C). The DiI positive fiber bundle of CFA were found posterior than RFA (Fig. 1C). Most of the callosal fibers from RFA and CFA were observed anterior to bregma (Fig. 1D). No DiI positive axonal fiber bundles were found in the posterior part of the corpus callosum (data not shown). These results suggest that in the rat brain, the motor components of the corpus callosum course mainly through the anterior parts of the corpus callosum.

**Figure 1.**
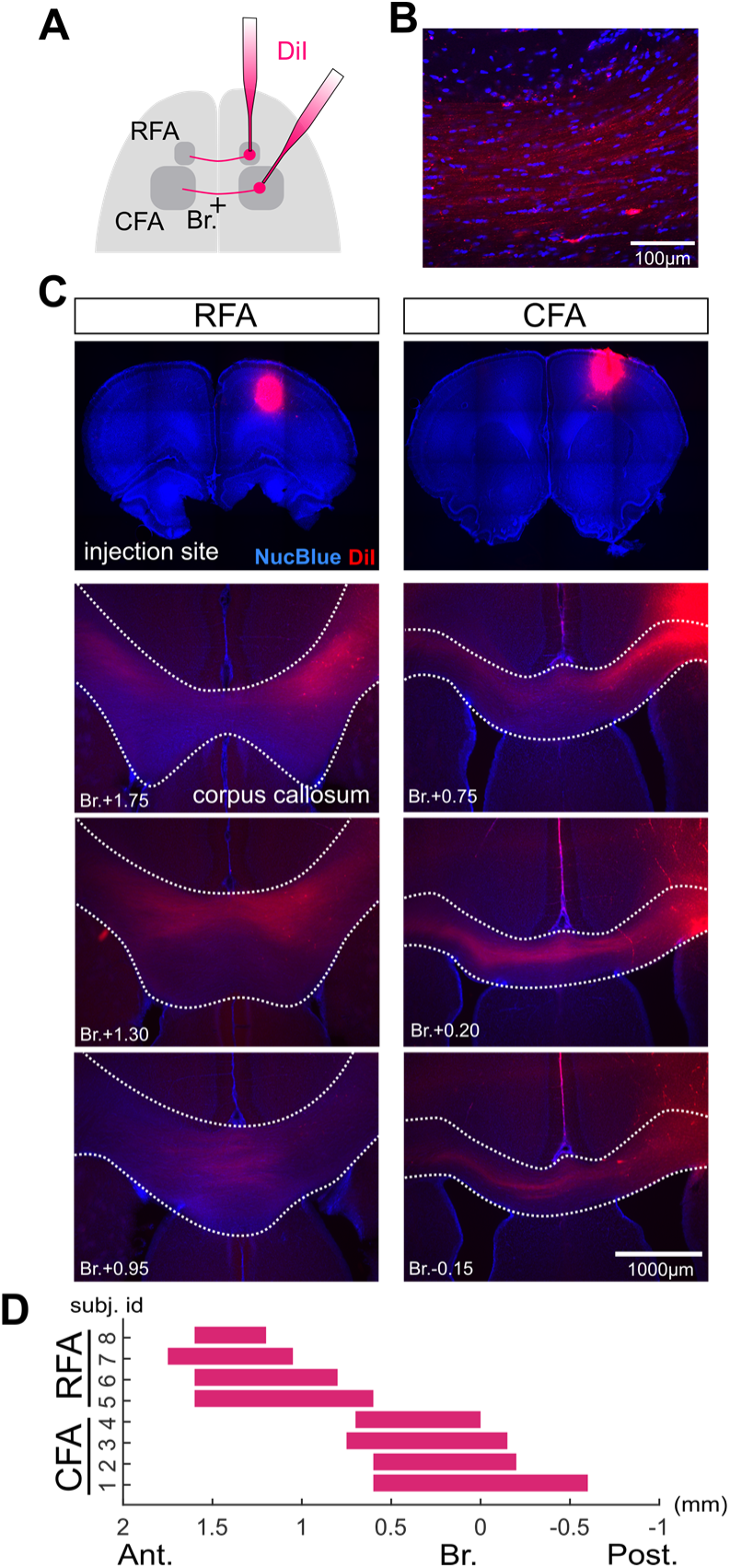
Motor fibers in anterior corpus callosum. (A) Schematic diagram of callosal axon fiber bundle labeling by 1,1’-dioctadecyl-3,3,3’,3’-tetramethylindocarbocyanine perchlorate (DiI). Axonal projections from two forelimb areas were labelled by an injection of DiI in either the rostral forelimb area or the caudal forelimb area. (B) Representative microimage showing DiI positive callosal fibers from CFA. (C) Anteroposterior profile of motor callosal path. Top row. Panels show the injection site in the RFA (left) and CFA (right). Rows 2-4 show the DiI positive callosal projection from RFA and CFA though the anterior part of the corpus callosum. (D) Summary of the anteroposterior location of axonal fiber bundle projection from CFA (n = 4, subject id 1, 2, 3, 4) and RFA (n = 4, subject id 5, 6, 7, 8).

### Local Lidocaine injection inactivates anterior corpus callosum

To study the causal role of neural signaling via the anterior corpus callosum, the local anesthetic Lidocaine was used to block axonal conduction in the corpus callosum. To validate the efficacy of Lidocaine *in vivo*, the efficacy of cortico-cortical axonal conductance was monitored by recording electrical stimulus-evoked population excitatory postsynaptic potential (pEPSP) responses. Intra-cortical current stimulation ICMS) was delivered in the right motor cortex while recording pEPSPs on the side contralateral to the ICMS (Fig. 2A, B). The latency of the pEPSP response was 13.13 ms (Std. Deviation = 2.03 ms), which is consistent with previous reports measured by antidromic spike (Wilson 1987; Soma *et al*. 2017), suggesting the pEPSP evoked by the ICMS is monosynaptic. Next, the current source density (CSD) profile was computed from the pEPSP traces. The CSD showed a significant sink (positive currents leaving extracellular medium) response at 1.05 mm from cortical pia (Std. Deviation = 0.30 mm). This shows the presence of an excitatory input from the contralateral cortex to cortical layers at a depth corresponding to layer 5. The ICMS-evoked sink response was maintained 30 min after saline injection (Fig. 2D). In contrast, 500nL of 2% Lidocaine injection significantly suppressed the sink amplitude (Fig. 2E). The suppression of the axonal conductance was observed from 10 min after injection and was sustained for 25 minutes (Fig. 2F). These result show that the 2% Lidocaine injection was effective in suppressing the cortico-cortical synaptic transmission. Thus, 2% Lidocaine was used to investigate the role of anterior corpus callosum in bimanual coordination in the behavioral experiments.

**Figure 2.**
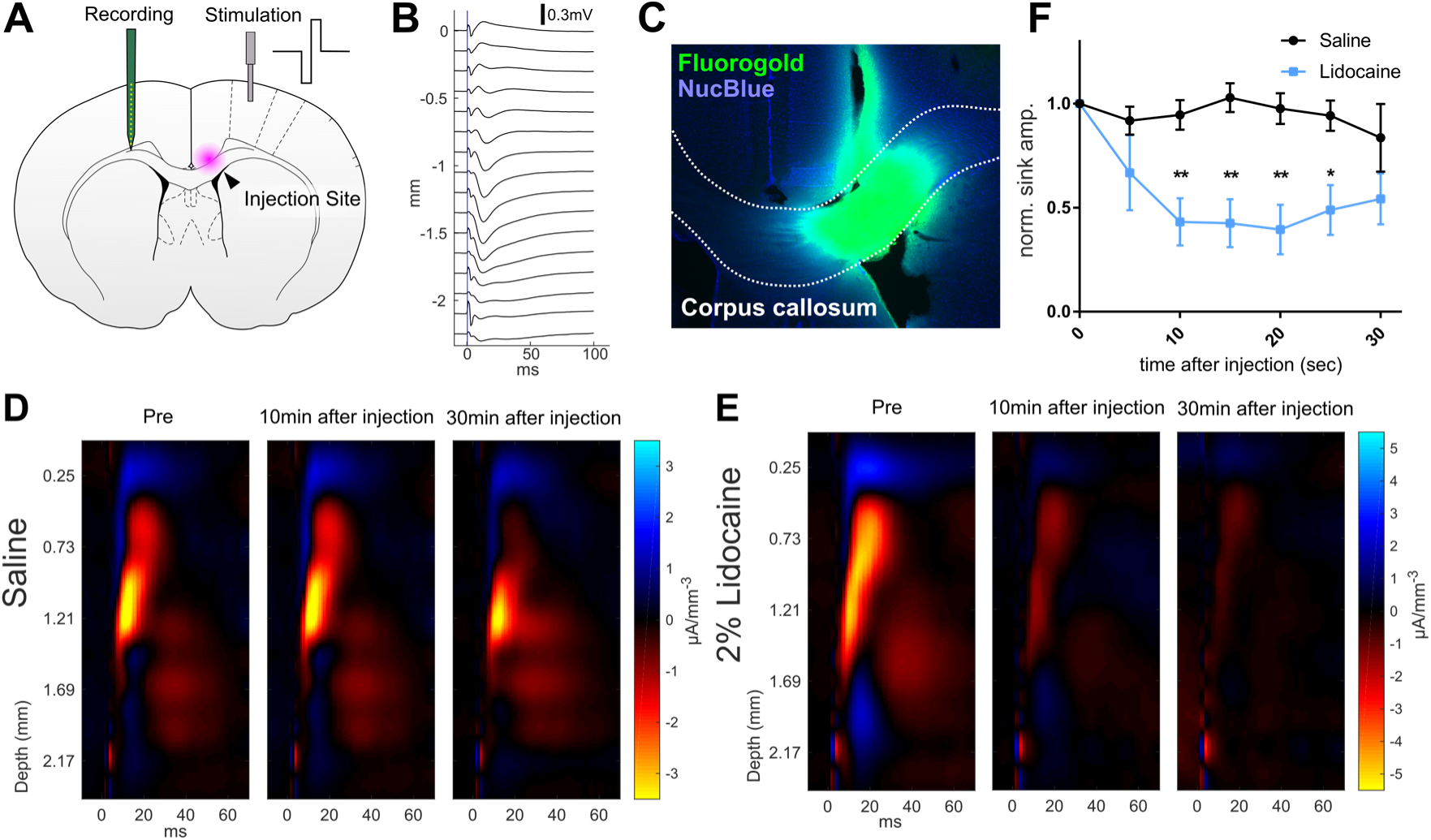
Lidocaine suppresses cortico-cortical signal transmission. (A) Schematic diagram of *in vivo* LFP recording. The stimulation electrode was placed in motor cortex, and the recording electrode was placed in the contralateral motor cortex. Lidocaine was locally injected into the anterior corpus callosum. (B) Mean pEPSPs at different cortical depths evoked by electrical stimulation (vertical blue line) of contralateral homotopic cortex. (C) Example of spread of solution from the injection site, visualized using 500 nL of FluoroGold (green) as an injected tracer. (D, E) Current source density (CSD) profiles corresponding to the recorded LFP across cortical depths. Negative current source was generated immediately after the onset of electrical stimulation. (D) Stimuli evoked CSD after saline injection to the corpus callosum. (E) Significant attenuation of stimulus evoked CSD response after Lidocaine injection. (F) Time course of normalized peak CSD sink response over 30 min period after Lidocaine injection. Significant attenuation of evoked responses was caused by the Lidocaine injection from 10 min to 25 min after injection (two-way ANOVA (F(1,6) = 22.57; p=0.0032) followed by Bonferroni’s multiple comparisons test (** p < 0.01, * p< 0.05). The error bars are ± SEM.

### Pharmacological inactivation of anterior corpus callosum in awake rats

Rats were trained to perform spontaneous food handling under head-fixed conditions (Fig. 3A). Rats consumed an annularly shaped food reward (Fig. 3B) by manipulating it in both hands, and the variety of forelimb movements during feeding were recorded via reflective markers attached to their wrists (Fig. 3A and C). The reconstructed forelimb trajectories in 3-D egocentric coordinate space were used for kinematic analysis of bimanual movements (Fig. 3D). Eleven successfully trained rats were subject to Lidocaine injections into the anterior corpus callosum. Each rat underwent test sessions consisting of three repeated daily cycles of Saline and Lidocaine injection conditions (Fig. 3E). After completion of all behavioral sessions, the locations of injection cannulae were confirmed by histology (Fig. 3F – H). Nine of eleven rats successfully received Lidocaine injection into the anterior corpus callosum (Table 1) and these rats were therefore used in the following data analyses.

**Figure 3.**
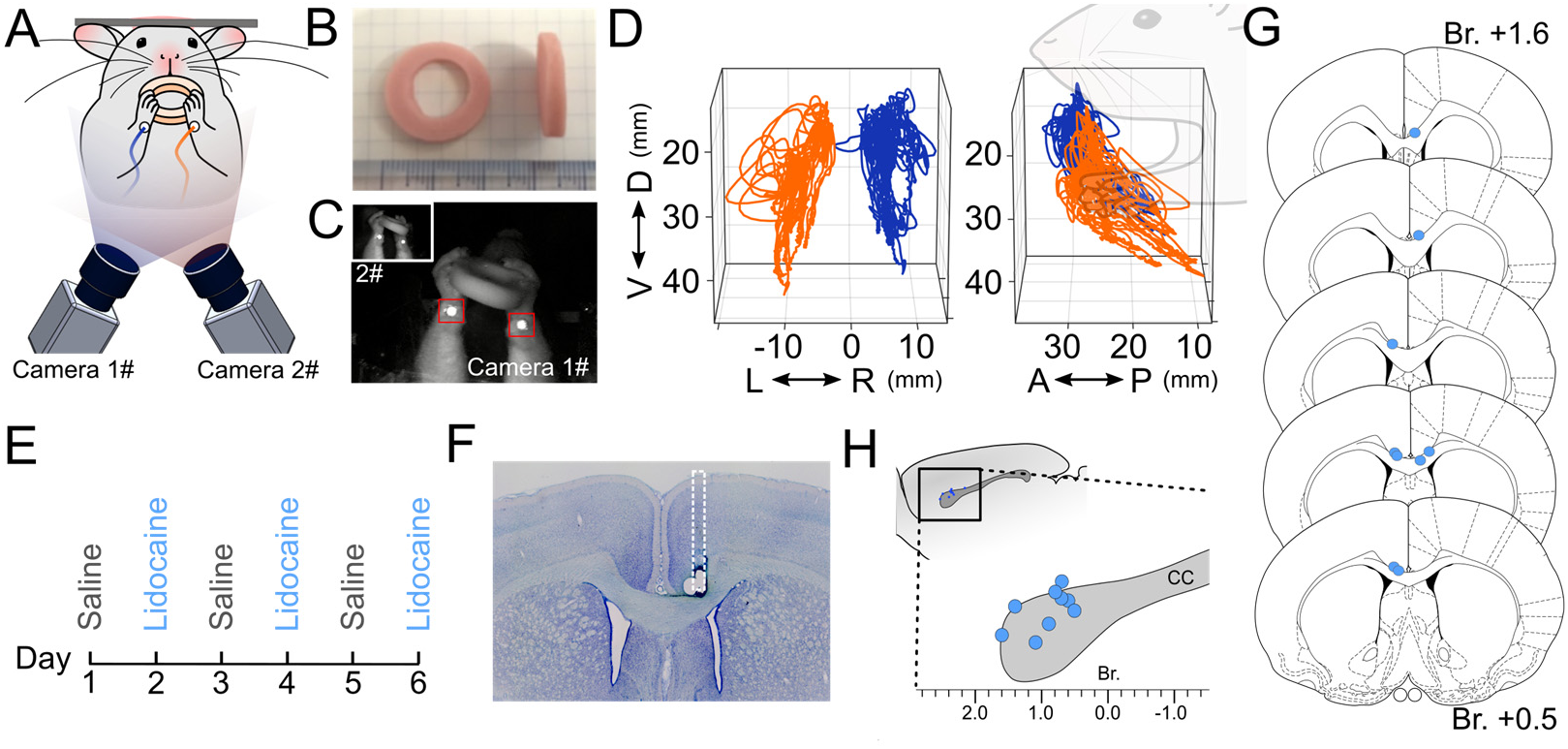
Pharmacological suppression of aCC by Lidocaine in awake rats. (A) Schematic diagram of recording system of forelimb motor behavior. Rats were placed in a head-fixing apparatus with transparent floor. (B) Food rewards were presented and consumed within the apparatus. (C) Forelimb motor behaviors were monitored during food consumption by two high-speed cameras placed below, with reflective markers attached to the lower side of each wrists. (D) Forelimb trajectories were traced by automatic detection of reflective markers and post-hoc 3-D reconstruction of the marker position. The positions of left (orange) and right (blue) forelimbs during food consumption are shown, viewed from the back (left panel) and side (right panel). Note the 3D trajectories were projected in a body centered (egocentric) coordinate frame. (E) Experiment schedule of behavioral recordings. Test sessions consisted of three repeated day-long cycles of conditions under baseline (Saline), and inhibition of anterior corpus callosum (Lidocaine) with more than 24 hr interval between days. (F) Verification of cannula implantation. The location of the tip of cannula was confirmed by Nissl staining. (G) Schematic of injection sites shown in coronal sections. (H) injection sites shown in sagittal plane. D, dorsal; V, ventral; A, anterior; P, posterior; R, right; L, left.

**Table 1.** Coordinates of injection sites

| Subject ID | Anterior from Bregma | Left from Bregma | Ventral from pia matter |
| --- | --- | --- | --- |
| Subj_38 | +0.8mm | +0.9mm | +3.0mm |
| Subj_39 | +0.9mm | +0.4mm | +3.5mm |
| Subj_40 | +1.1mm | -0.9mm | +3.5mm |
| Subj_41 | +1.8mm* | +0.5mm | +3.0mm |
| Subj_42 | +1.6mm | +0.6mm | +3.0mm |
| Subj_43 | +1.8mm* | +0.7mm | +3.0mm |
| Subj_47 | +0.7mm | -0.9mm | +2.8mm |
| Subj_49 | +0.5mm | -0.8mm | +3.2mm |
| Subj_50 | +0.7mm | -0.8mm | +3.2mm |
| Subj_51 | +1.4mm | +0.5mm | +3.0mm |
| Subj_52 | -0.4mm* | +0.8mm | +3.0mm |
| Subj_53 | +0.6mm | -1.0mm | +3.2mm |

\*missed target

### No effect of anterior corpus callosum blockade on proportion of bilateral movements

We first examined whether the blockade of the anterior corpus callosum affects the proportion of bilateral versus unilateral movement, based on the ratio of the faster to slower forelimb speeds (Fig. 4A). Bilateral movements were defined as those in which the slower forelimb moved with at least half the speed for the faster limb (SR ≥ 0.5, Fig. 4B), and unilateral movements were defined as those in which the slower limb moved at less than half the faster limb (SR < 0.5, Fig. 4C) (Igarashi and Wickens 2019). The speed ratio was computed along with trajectories of forelimbs for all recorded trials of two conditions (Saline and Lidocaine, Fig.4D). As seen in the colored trajectories, the most time was spent in bilateral movement (Fig. 4D). Quantitative analysis revealed that in the saline control group, 89.37 % of time movements were classified as bilateral (Fig. 4E), while the remaining 10.63% were unilateral movements (Fig. 4F). After injection of Lidocaine into the anterior corpus callosum, neither reduction nor increase was observed (Fig. 4E,F). These results suggest that the anterior corpus callosum does not mediate the balance of movement speed between two forelimbs during feeding behavior.

**Figure 4.**
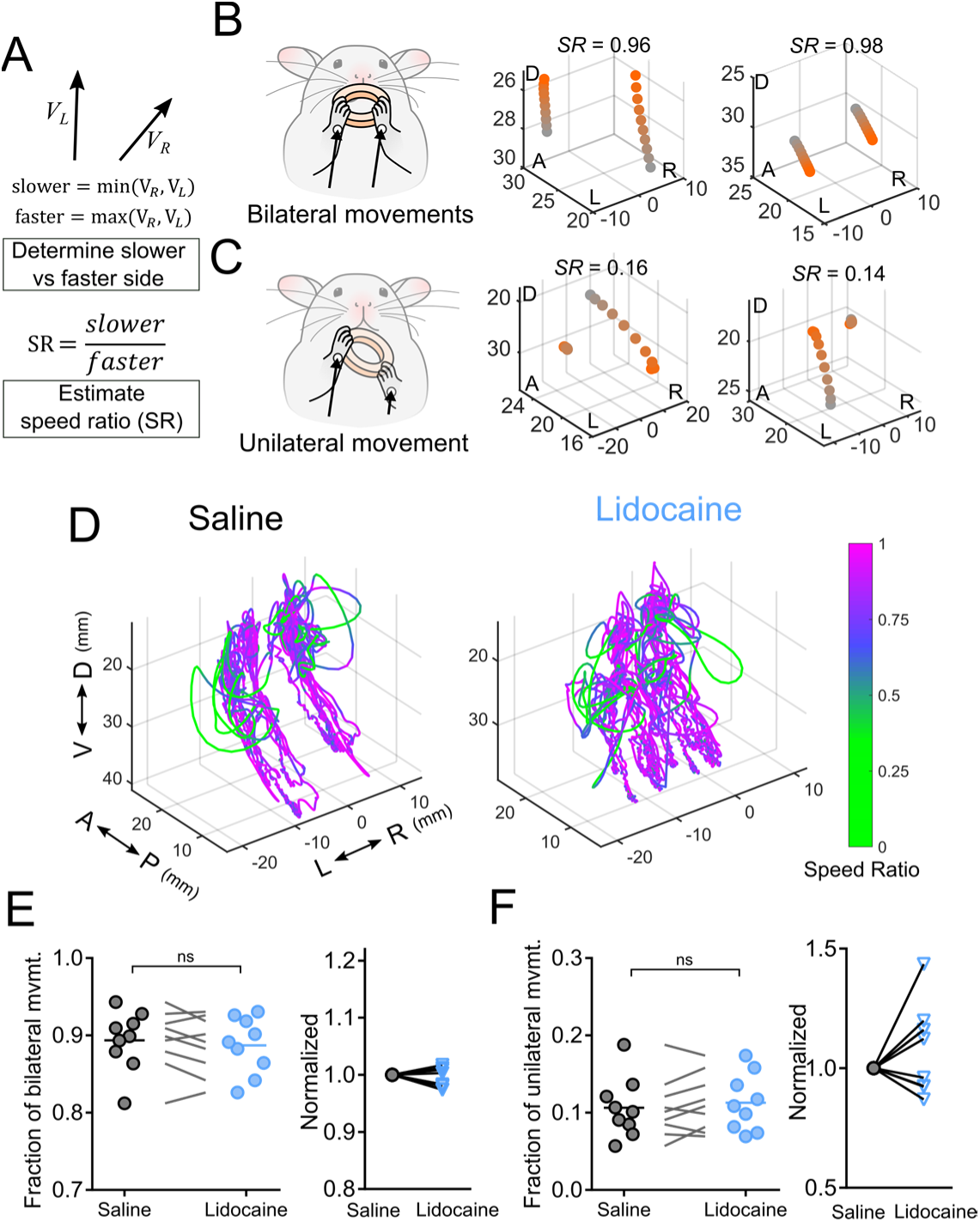
Anterior corpus callosum does not mediate the balance of movement speed between forelimbs during feeding behavior. (A-C) Kinematic data were analyzed by speed ratio function to calculate balance in movement speed between left and right forelimb. (A) Speed ratio computed the ratio of faster and slower forelimb speeds. Higher speed ratio indicates the speed closer in two forelimb. (B) Schematic drawing of bilateral forelimb movements (left panel). The displacements of forelimb are not significantly different across two forelimbs. Representative example of forelimb trajectory segment in bilateral mode showing SR ≥ 0.5 (middle and right panel). (C) Unilateral forelimb movements (left panel). The displacement of one side is significantly greater than the contralateral forelimb. Two representative examples of forelimb trajectory segment in unilateral mode showing SR < 0.5 (middle and right panel). (D) Forelimb trajectories of saline (left) and Lidocaine (right) injected group. The speed ratio was overlain on the 3D trajectories in color scale. (E-F) Time fraction on unilateral and bilateral forelimb movements were unchanged. (E) Quantitative analysis of the fraction of time spent on bilateral movements (SR ≥ 0.5) during food consumption (left), and the normalized change (right) (paired t-test, p = 0.2326, n = 9). (F) Quantitative analysis of the fraction of time spent on unilateral movements (SR < 0.5) during food consumption (left), and the normalized change (right) (paired t-test, p = 0.2326, n = 9). D, dorsal; V, ventral; A, anterior; P, posterior; R, right; L, left. Numbers in 3-D plots are expressed in millimeters.

### Anterior corpus callosum plays role in symmetric forelimb movements

Next, we tested the effect of the attenuation of anterior corpus callosum on symmetry in movement direction during spontaneous food manipulation. To measure symmetry in movement direction, the asymmetry index was used (Igarashi and Wickens 2019). The velocity of left forelimb and the mirrored right forelimb was used to compute the asymmetry index, represented as the angle θ between the movement vectors of the forelimbs (Fig. 5A). The asymmetry index values were calculated over the 3D trajectories (Fig. 5D). Quantitative analysis revealed that in saline control group 57.17 % of the time movements were classified as symmetric (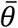 ≤ *π*/4, Fig. 4E), and the remaining 42.83% were classified as asymmetric (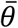 > *π*/4, Fig. 4F), which was consistent with our previous report (Igarashi and Wickens 2019). After injection of Lidocaine into the anterior corpus callosum, a reduction of symmetric movements was observed. The fraction of time in the symmetric mode decreased to 53.87 % (Fig. 4E), and in turn, asymmetric movement increased (46.13 %, Fig. 4E). The finding of reduction of symmetric movements was robust in the face of variations in the parameters of the sliding time window used for segmentation (supplemental figure 2). The reduction of symmetric movements was not observed in a naïve control group (supplemental figure 3). These results suggest that the anterior corpus callosum plays an important role in symmetric bilateral forelimb movements during spontaneous food handling behavior.

**Figure 5.**
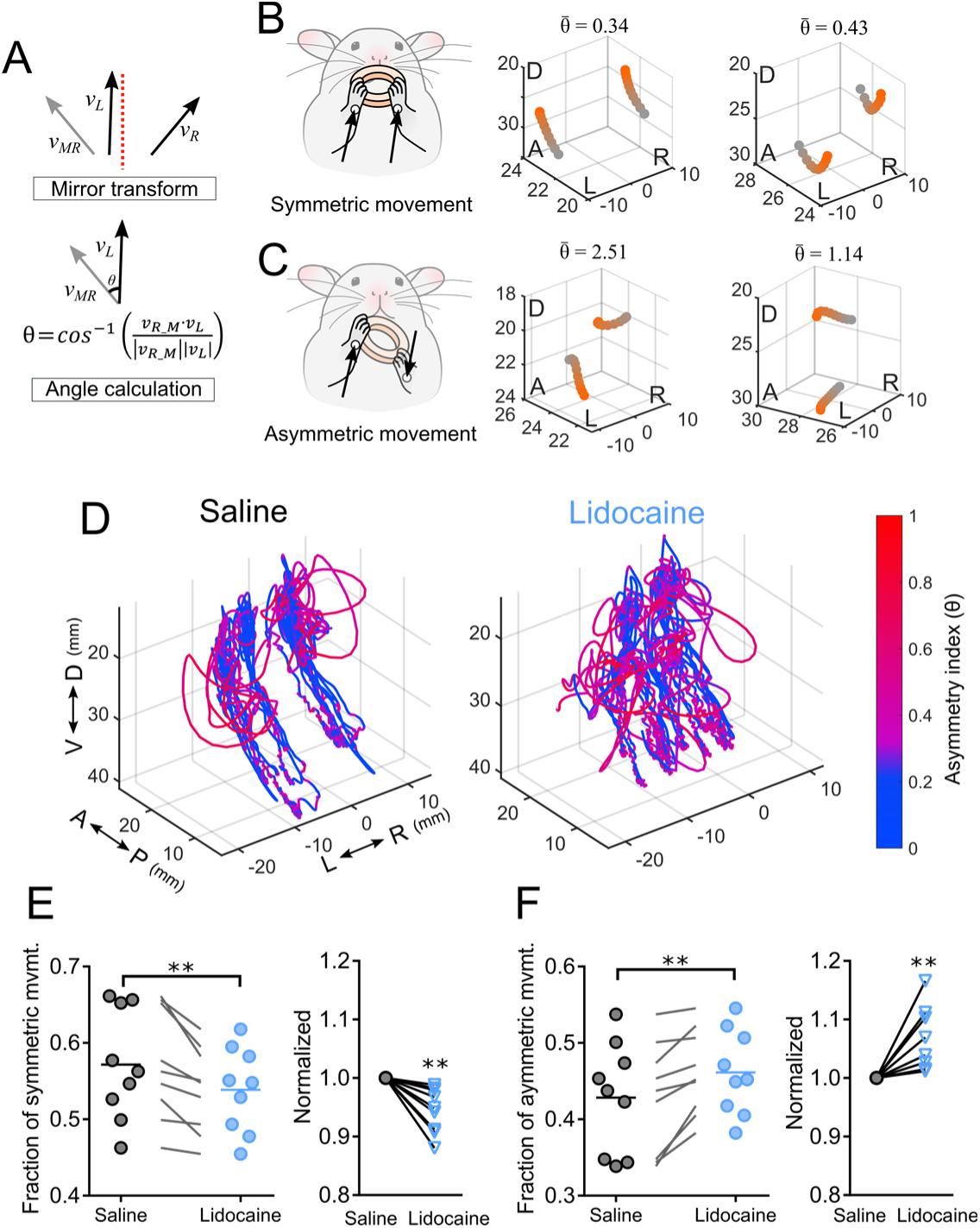
Blockade of anterior corpus callosum made the symmetric forelimb movements more asymmetric in movement direction. (A) Asymmetry index was computed by the difference of movement direction between the left forelimb velocity *υ*_L_ and the mirrored right forelimb velocity *υ*_MR_. The mirror transformation was applied to the velocity vector *υ*_R_ to create the mirrored right forelimb velocity vector *υ*_MR_. The asymmetry index was then computed by the angle between *υ*_L_ and *υ*_MR_. Asymmetry index represents the degree of divergence in movement direction. (B) Schematic drawing of symmetric forelimb movements (left panel). The movement directions are not significantly different across forelimbs in symmetric movements. Representative fraction of forelimb trajectories in symmetric mode showing 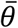 ≤ *π*/4 (middle and right panel). (C) Schematic drawing of asymmetric forelimb movements (left panel), where the movement direction diverges. Two representative forelimb trajectories in asymmetric mode are shown 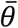 > *π*/4 (middle and right panel). (D) Colored forelimb trajectories of saline (left) and Lidocaine injected group (right). The asymmetry index was indicated using a color scale. (E-F) Reduction of the ratio of symmetric movements and the increase of asymmetric movements. (E) Quantitative analysis of the fraction of time spent on symmetric movements 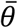 ≤ *π*/4 during food consumption (left), and the normalized change (right) (paired t-test, p = 0.0043, n = 9). (F) Quantitative analysis of the fraction of time spent on asymmetric movements 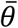 > *π*/4 during food consumption (left), and the normalized change (right) (paired t-test, p = 0.0043, n = 9). Note the fraction of time of asymmetric movements complements symmetric movements. D, dorsal; V, ventral; A, anterior; P, posterior; R, right; L, left. Numbers in 3-D plots are expressed in millimeters.

### Effect of anterior corpus callosum blockade on global motor function

The suppression of anterior corpus collusum neurotransmission had no significant effect on task performance or measures such as mean consumption time and success rate (supplemental figure 1A,B). There was also no effect of variation in injection location on mean forelimb movement speed, suggesting that the Lidocaine injection did not cause motor disability of forelimb by spreading to the motor cortex adjacent to the injection (supplemental figure 1C,D).

## Discussion

The role of corpus callosum in bimanual coordination was examined by analyzing forelimb kinematics during object manipulation, before and after pharmacological suppression of the anterior corpus callosum. Neural tracing showed that the fiber bundle from the forelimb motor areas in rats passes through the anterior portion of the corpus callosum. We confirmed with electrophysiology that neural transmission in the anterior transcortical pathway was attenuated by injections of local anesthetic, which reduced the LFP response evoked by ICMS of the contralateral hemisphere. Kinematics of forelimb movements with and without suppression of the anterior corpus callosum were then compared during bimanual food handling. Suppression of the anterior corpus callosum decreased the fraction of forelimb movements that were bilaterally symmetric, whereas the balance of movements speed timing and other global scores were unchanged. These results suggest that the anterior corpus callosum contributes to symmetry in the form of bilateral forelimb movements. To our knowledge, this is the first study to investigate the role of the rodent motor corpus callosum in bimanual coordination, extending previous knowledge of the role of corpus callosum in bimanual coordination in daily feeding behavior.

Studies of the location of motor fibers in the corpus callosum of human, using diffusion tensor imaging, have shown that the callosal motor fibers are found from the anterior part (genu) to the posterior body and isthmus of the corpus callosum (de Lacoste *et al*. 1985; Hofer and Frahm 2006; Wahl *et al*. 2007; Gooijers and Swinnen 2014). In the present study, the location of callosal motor fibers of rats was studied using a neural tracer. The results suggest that the callosal motor fibers of rats are most dense in the anterior part of the corpus callosum. The RFA callosal motor fibers ran more anteriorly than the CFA callosal motor fibers. This result is consistent with the projection map of the *Allen Mouse Brain Connectivity Atlas* (http://connectivity.brain-map.org, Oh *et al*., 2014), which shows that mouse motor areas are connected by the anterior part of corpus callosum. Therefore, the anterior corpus callosum was targeted in the present study.

In order to reversibly block axonal conduction of the corpus callosum, without altering the neurons of origin, lidocaine was injected into the anterior corpus callosum. The blockade of axonal conduction was confirmed by ICMS and LFP recordings. Control records showed that a significant sink response across cortical laminar in the homotopic cortex contralateral to ICMS, with a maximum at a level corresponding to layer 5, which is consistent with previous report in the sensorimotor cortex of rats (Chapman *et al*. 1998). Since 99% of the axonal fibers of the corpus callosum are excitatory fibers which originate from glutamatergic intratelencephalic neurons (Shepherd 2013; Harris and Shepherd 2015), the sink responses in the present study almost certainly reflects excitatory inputs from the contralateral cortex to these layers. There remains, however, a theoretical possibility that the evoked sink might be caused by polysynaptic events such as peripheral inhibitory inputs mediated by the callosal-interneuron pathway (Palmer *et al*. 2012; Kokinovic and Medini 2018), or other indirect components via posterior part of cerebral cortices. We found the latency of the response is consistent with previous reports measured by antidromic spike (Wilson 1987; Soma *et al*. 2017), suggesting that the sink observed in the present study is indeed caused by monosynaptic excitatory synaptic inputs. Injecting lidocaine into the anterior corpus callosum significantly suppressed the sink of LFP, confirming that axonal conduction was blocked in the direct excitatory interhemispheric connection between homotopic motor cortical areas. Taken together, we suggest that the lidocaine lesion effectively attenuates the excitatory interhemispheric connections.

In the present study, pharmacological blockade of axonal conduction was used instead of corpus callosotomy, which has been a widely in previous studies of the role of interhemispheric communication in rodents (Mohn and Russell 1981; Noonan and Axelrod 1992; Sullivan *et al*. 1993; Li *et al*. 2016) and non-human primates (Mark and Sperry 1968). In human, patients received callosotomy as treatment of seizure have been subject to test the role of corpus callosum in visual perception (Gazzaniga *et al*. 1962), and motor control (Franz *et al*. 1996; Ivry and Hazeltine 1999; Franz *et al*. 2000; Kennerley *et al*. 2002). The pharmacological blockade has the advantage of being reversible. In the present study two rats showed recovery from suppression approximately 30 min after the Lidocaine injection, which provided rapid reversibility of the pharmacological suppression of the corpus callosum. For studies requiring longer duration of suppression, longer-acting sodium channel blockers, such as QX-314, are available (Binshtok *et al*. 2009).

In the present study, inhibition of aCC did not alter the ratio of bilateral to unilateral movement during food handling, suggesting the balance of movement speed between two forelimbs was not changed. During food handling, the speed forelimb movements were well-synchronized, which is represented as predomination of bilateral movements, as previously reported (Igarashi and Wickens 2018). The predomination of bilateral synchronization has been reported in the studies done by human. For example, in human, synchrony of movement timing can be observed in rhythmic bimanual finger tapping; and the in-phase mode (simultaneous finger tapping with no phase shift; ∅ = 0°) is more stable than anti-phase mode (tapping with alternation; ∅ = 180°) (Yamanishi *et al*. 1980; Schoner and Kelso 1988). Interestingly, the bilateral synchrony of temporal coupling was well-preserved in callosotomy patients (Tuller and Kelso 1989; Ivry and Hazeltine 1999). Donchin *et al*. (1999) proposed the involvement of a subcortical structure, a central pattern generator (CPG), in synchronizing bimanual movements. This proposal might explain how conserved timing synchrony is possible without corpus callosum, because this would leave a subcortical CPG intact. However, there remain unknown whether the predomination of bilateral movements during spontaneous feeding in rodents and the timing synchronization during rhythmic bimanual task in human are mediated by the similar neural substrates. Further investigation is needed to understand how rodents achieve bilateral control of movements speed in a highly balanced manner without motor-motor interhemispheric connection.

In contrast, inhibition of aCC by Lidocaine reduced the symmetry of bilateral forelimb movements. A possible interpretation of this reduction of movement symmetry is that the corpus callosum is necessary for the symmetry of bimanual movements. Humans exhibit a tendency toward spatially symmetric movements during bimanual tasks (Franz 1997; Swinnen *et al*. 1998; Walter *et al*. 2001). There is evidence that this depends on the corpus callosum. Spatial coupling in a bimanual drawing task (drawing of different forms by two hands) was reduced in split-brain patients (Franz *et al*. 1996). In addition, disruption of temporal synchrony in split-brain patient became evident when the task involved spatial requirements in addition to timing (e.g., continuously drawing circle in a 2-D plane) (Kennerley *et al*. 2002), suggesting that the symmetric form of bilateral forelimb movements is mediated by corpus callosum in humans, consistent with the present results obtained in rats during natural eating behavior. In rats, the present study suggests that the frequent symmetrical upward and downward bimanual reaching action that occurs in feeding may be mediated by the anterior corpus callosum. However, it still remains unclear whether the symmetric movements mediated by the corpus callosum are responsible for the specific motor pattern of bimanual acts, such as upward bimanual reaching during food-to-mouth behavior, and downward bimanual reaching during tearing of food. Further work is needed to address this issue.

The mechanism underlying the contribution of the corpus callosum to symmetric bimanual movements is not well understood at the cellular level. It has been unclear whether the connection is functionally excitatory or inhibitory. For example, the attenuation of the spread of seizure and loss of information integration in split-brain patients suggests that the corpus callosum is used for excitatory interhemispheric signal transmission (Gazzaniga *et al*. 1962; Spencer *et al*. 1988; Fuiks *et al*. 1991). Repeated stimulation to corpus callosum (kindling stimulation) forms bilateral representation in rat forelimb motor area supporting the excitatory role of corpus callosum (Teskey *et al*. 2002). The purpose of such a connection in motor function is not yet known. One possibility is suggested by the proposal of Li *et al*. (2016), in the form of a modular attractor model comprised of independent modules encoding particular actions. In the model, callosal excitatory connections link homotopic modules in each hemisphere. Given the existence of topographical maps related to forelimb movement form (Young *et al*. 2011; Brown and Teskey 2014) and direction (Hira *et al*. 2015), linking these across hemispheres might contribute to spatially symmetric movement. The effect of the lesion on the extent of symmetric movement might then make sense if these excitatory callosal projections are connected to functionally homotopic areas across motor cortices.

On the other hands, it has been postulated that the corpus callosum inhibits neurons in the contralateral hemisphere (interhemispheric inhibition, IHI) (Ferbert *et al*. 1992; Hubers *et al*. 2008). At the cellular level, IHI would result from dysynaptic connections involving inhibitory interneuron activated by excitatory inputs of the corpus callosum (Palmer *et al*. 2012; Kokinovic and Medini 2018). The electrical stimulation of corpus callosum causes EPSP in the sensorimotor cortex followed by IPSP (Chapman *et al*. 1998; Teskey *et al*. 1999). The functional significance of such inhibition is suggested by experiments in which cooling of contralateral somatosensory cortex unmasked larger receptive field, therefore, less selective to sensory inputs (Clarey *et al*. 1996). A similar report has been reported in rats (Pluto *et al*. 2005). These are consistent with the idea that the callosal connection provides a source of inhibition for shaping the finer receptive field. It should be noted, however, that the excitatory model and inhibitory model are not mutually exclusive. Rather both excitatory and inhibitory connections might play important roles in activating the contralateral motor cortex to perform finer bilateral forelimb movements accurately by inactivating unnecessary movements.

The present kinematic analysis revealed a reduction of symmetric bimanual movements by blockade of aCC, however, the rats were still able to perform the food handling under head-fixation without significant reduction in mean consumption time and successful completion rate. This raises the question of why the blockage of the aCC did not significantly modify bimanual eating behavior. As described above, prior studies in primates demonstrated that the corpus callosum plays an important role in bimanual coordination in a variety of tasks. However, it has been reported that callosotomy patients had less difficulty in familiar bimanual actions such as tying a shoe and opening drawer (Franz *et al*. 2000; Serrien *et al*. 2001). Therefore, it is possible that the corpus callosum is not crucial for the execution of familiar bimanual actions such as food handling. Nevertheless, information exchange between the two sides of the body is necessary for highly synchronized movement direction, and the present study suggests that this is mediated by the aCC.

## Author contributions

M.I. and J.W. designed experiments. M.I. conducted experiments and drafted manuscript and figures. Y.A. conducted histology. J.W. revised manuscript.

## Acknowledgment

This work was supported by Japan Society for the Propotion of Science Reserch Fellow Grant-in-Aid 16J05329.

---

**Supplemental figure 1.**
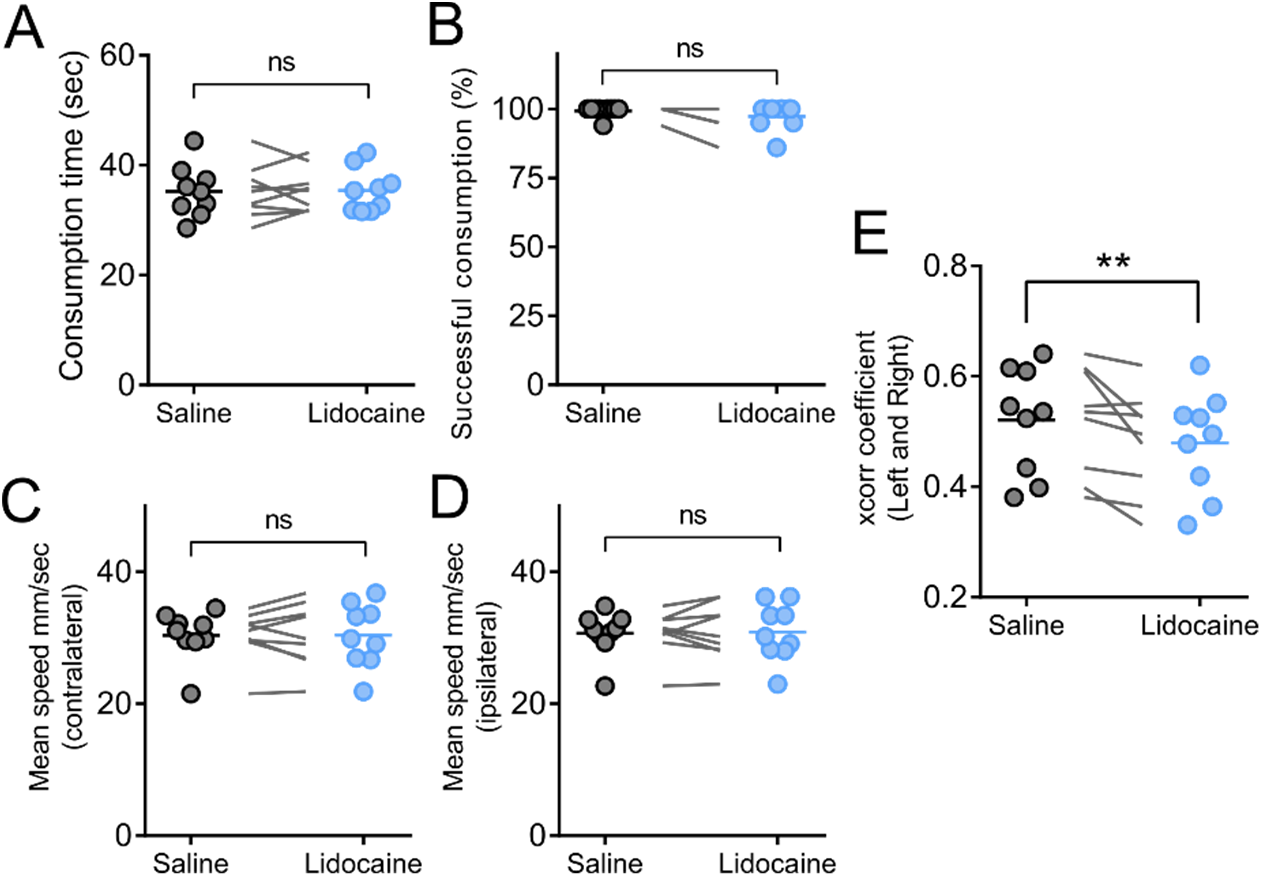
Inhibition of aCC did not alter global motor/task performances but cross correlation. (A) Mean consumption time (ns, p=0.8635). (B) Successful rate was unchanged (ns, p=0.0909). (C) The mean speed of forelimb contralateral to the Lidocaine infusion (ns, p=0.9825), and the ipsilateral (D, ns, p =0.8361). (E) cross-correlation between velocity of left and mirrored-right forelimb (significant, p=0.0255). All statistical significance was validated by paired t-test (n=9).

**Supplemental figure 2.**
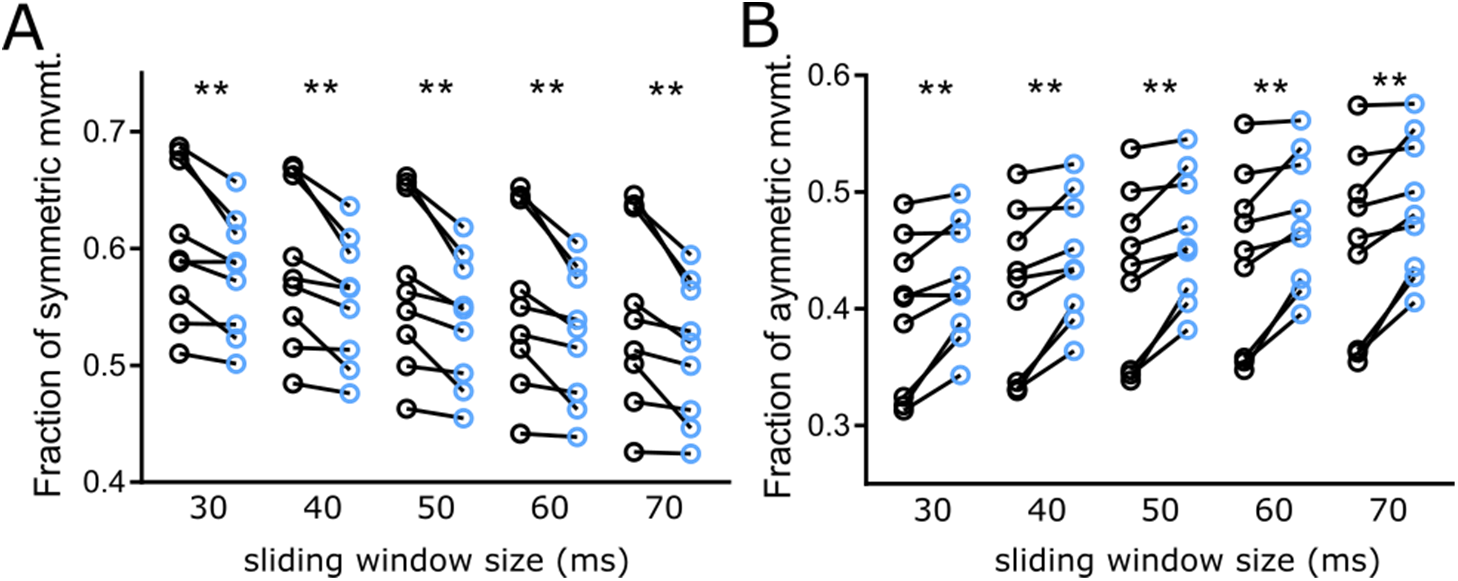
Reduction of symmetric movements with different sliding window sizes. To test the effect of size of sliding window in the kinematic analysis, five different time windows were used to reanalyze symmetric and asymmetric ratio. The reduction of symmetric fraction and increase of asymmetric fraction was consistently observed all different time window.

**Supplemental figure 3.**
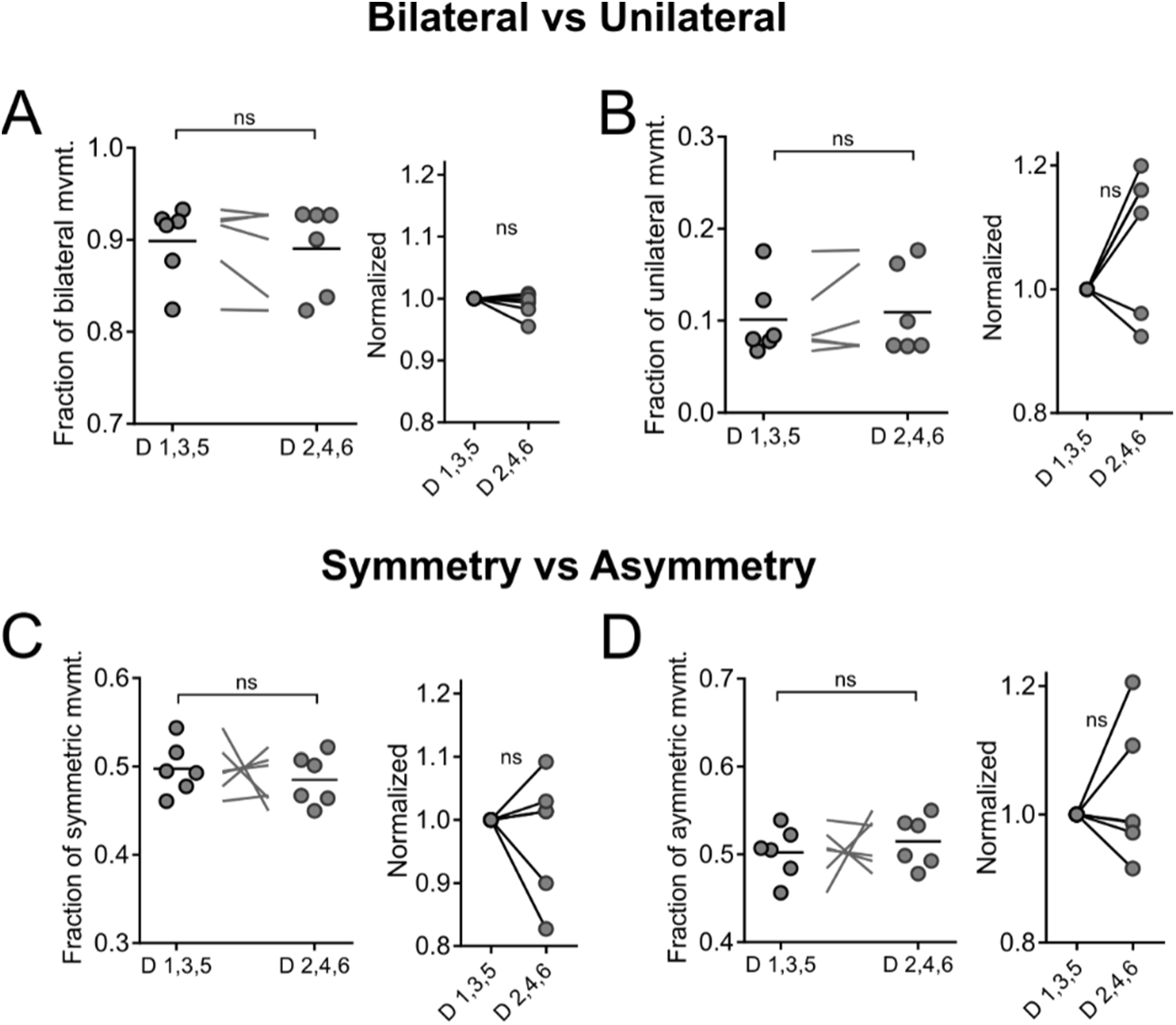
No significant difference in Naïve control group. (A) Quantitative analysis of the fraction of time on bilateral movements SR ≥ 0.5 during food consumption (left), and the normalized value (right) (paired t-test, p = 0.293, n = 6). The horizontal bar in the left panel is mean. (B) Quantitative analysis of the fraction of time on unilateral movements SR < 0.5 during food consumption (left), and the normalized value (right) (paired t-test, p = 0.293, n = 6). (C) Quantitative analysis of the fraction of time on symmetric movements 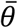 ≤ *π*/4 (left), and the normalized value (right) (paired t-test, p = 0.574, n = 6). (D) Quantitative analysis of the fraction of time on asymmetric movements 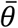 > *π*/4 (left), and the normalized value (right) (paired t-test, p = 0.574, n = 6). Note the fraction of time of asymmetry and unilateral movements are counterpart of symmetric movements and bilateral movements respectively.

